# NanoR: a user-friendly R package to analyze and compare nanopore sequencing data

**DOI:** 10.1101/514232

**Authors:** Davide Bolognini, Niccolò Bartalucci, Alessandra Mingrino, Alessandro Maria Vannucchi, Alberto Magi

**Affiliations:** Department of Experimental and Clinical Medicine, University of Florence, Largo Brambilla 3, Florence, Italy

## Abstract

MinION and GridION X5 from Oxford Nanopore Technologies are devices for real-time DNA and RNA sequencing. On the one hand, MinION is the only real-time, low cost and portable sequencing device and, thanks to its unique properties, is becoming more and more popular among biologists; on the other, GridION X5, mainly for its costs, is less widespread but highly suitable for researchers with large sequencing projects. Despite the fact that Oxford Nanopore Technologies’ devices have been increasingly used in the last few years, there is a lack of high-performing and user-friendly tools to handle the data outputted by both MinION and GridION X5 platforms. Here we present NanoR, a cross-platform R package designed with the purpose to simplify and improve nanopore data visualization. Indeed, NanoR is built on few functions but overcomes the capabilities of existing tools to extract meaningful informations from MinION sequencing data; in addition, as exclusive features, NanoR can deal with GridION X5 sequencing outputs and allows comparison of both MinION and GridION X5 sequencing data in one command. NanoR is released as free package for R at https://github.com/davidebolo1993/NanoR.

## Introduction

During the last decade, second-generation sequencing technologies have definitely improved our ability to detect genetic variants in any organism. However, the short reads (100-500 base pairs) generated by these platforms do not allow to detect complex structural variants such as copy number variations [1]. In the last few years, third-generation sequencing (TGS) technologies have risen to prominence; Oxford Nanopore Technologies (ONT) and Pacific Biosciences [2] allow single-molecule and long-read sequencing with different strategies: while Pacific Biosciences uses a single-molecule real-time approach, ONT’s devices (MinION, GridION X5 and PromethION) analyze the transit of each DNA molecule through a nanoscopic pore and measure its effect on an electric current.

Among ONT’s devices, MinION and GridION X5 are noteworthies.

MinION is the only low-cost (commercially available by paying a starter-pack fee of $1,000), portable (~ 10 cm in length and weighs under 100 g) and real-time DNA and RNA sequencer: each MinION Flow Cell, which contains the proprietary sensor array Application-Specific Integrated Circuit (ASIC) and 2048 R9.4 (for sequencing kits 1D) or R9.5 (for sequencing kits 1D^2^) nanopores arranged in 512 channels, is capable to generate 10–20 Gb of DNA sequence data over a 48 hours sequencing run [3]. For its unique properties, from the launch of MinION Access Programme (April 2014), MinION has become increasingly popular among biologists.

Although it is definitely more expensive (~$50,000.00), GridION X5 fills the gap between MinION and ONT’s top-of-the-range PromethION. Indeed, GridION X5 is a small benchtop sequencer that allows to use up to five MinION Flow Cells at the same time; as result, GridION X5 drastically increases the ouput of a single MinION experiment to more than 100 Gb of DNA sequence data over a 48 hours sequencing run, making this instrument suitable for researchers with broader scope projects.

While library preparation is fast and easy when working with ONT’s devices, the biggest problem researchers have to face when dealing with these TGS technologies is certainly the analysis of their outputs.

The main output of ONT platforms are binary *hdf5* files [4] organized in a hierarchical structure with groups, datasets and attributes and stored using the *.fast5* extension. After MinION release 18.12 and GridION X5 release 18.12.1, *.fast5* files generated by this platforms are basecalled using the GPU-enhanced basecaller Guppy, which is more than an order of magnitude faster than previous CPU-based basecalling softwares, and stored into passed and failed folders according to their quality: passed *.fast5* files have mean quality ≥ 7 while failed *.fast5* files have quality < 7. Moreover, even if MinION and GridION X5 produce by default a single *.fast5* file per read (single-read *.fast5* files), since ONT sequencing experiments frequently generate millions of reads, a new multi-read *.fast5* files format has been introduced, which is more practical for data transfer and data querying considerations. For each sequencing experiment, in addition, a sequencing summary *.txt* file and *.fastq* files, splitted into passed (*i.e. .fastq* files and failed (*i.e. .fastq* files according to the aforementioned quality creteria, are generated.

As for the other next-generation sequencing platforms, data generated by ONT’s devices must be rapidly assessed in order to determine how useful the data may be in making biological discoveries as higher the quality of data is, more confident the conclusions are.

In the past years a number of tools designed specifically for MinION single-read *.fast5* files visualization became available, including poretools [5], poRe [6] and HPG pore [7].

Poretools was the first tool released for MinION data analysis and it implements several functions to compute statistics and extract key informations (*i.e.* metadata) both from entire .fast5 files datasets and single .fast5 files; however, most of poretools’ functions designed to return an overview of the sequencing experiment need to be run on the entire .fast5 file dataset, which basically means reading the same dataset multiple times to extract different informations, causing a waste of time when the user need to call multiple functions. Poretools implements also a function to return a metadata table from a dataset of .fast5 files in a single run, but does not provide any function to compute/plot statistics from this table.

PoRe shares most of poretools’ features and in 2017 a GUI for parallel and real-time extraction of metadata from MinION reads has been released [8]: by using multiple cores, the GUI version makes it faster to extract informations from .fast5 files but, again, its downstream statistical analysis is extremely limited.

HPG pore, at last, features the capability to extract metadata from .fast5 files and plot some statistics by parallelizing the job inside an Hadoop distributed file system, which lets you store and handle massive amount of data on a cloud of machines: for this reasons, HPG pore’s true potentiality can be exploited only in a Hadoop cluster. In addition, HPG pore can only be run on Linux machines.

Taking all this into account, we developed NanoR, an up-to-date package for the cross-platform statistical language and environment R [9], which is growing more and more popular with researchers, to analyze and compare both MinION and GridION X5 1D sequencing data. Our package, which has been tested on all the three main computer operating systems (Linux, MacOSX and Windows), is designed to be extremely easy to use and allows users to retrive a complete overview of their sequencing runs within acceptable time frames, offering the possibility to parallelize the extraction of metadata from .fast5 files, which is the most time-consuming step of the analysis process.

## Results

NanoR is a cross-platform package for the statistical language and environment R, requiring the following additional packages:

- ggplot2 [10]
- reshape2 [11]
- scales [12]
- RColorBrewer [13]
- gridExtra [14]
- rhdf5 [15]
- ShortRead [16]

Further informations on the versions of the aforementioned packages needed to run NanoR are available in the *Availability and requirements* section.

NanoR is built on 4 main functions:

- *NanoPrepare()*: prepares MinION/GridION X5 data for the following analyses
- *NanoTable()*: generates a table that contains metadata for every sequenced read
- *NanoStats()*: plots statistics for the sequencing run
- *NanoFastq()*: extract/filter *.fastq* sequences from MinION/GridION X5 data

Each of these functions exists both in a “M” version and in a “G” version (*e.g.* for *NanoPrepare()* exists both the *NanoPrepareM()* and the *NanoPrepareG()* version). This identifiers refer to the original behaviour of MinION and GridION X5. Indeed, on the one hand, MinION originally outputted only basecalled *.fast5* files from which metadata could be extracted in order to generate an overview of the sequencing run; on the other, GridION X5 outputted *.fast5* files without basecalling informations added, causing the need to use the simultaneosuly generated sequencing summary files to get a thorough analysis of the experimental run. After MinION release 18.12 and GridION X5 release 18.12.1, the format of the data generated by the aforementioned platforms have been standardized. Using the “M”/“G” notation, NanoR offers the possibility to choose if the analysis will be done starting from basecalled *.fast5* files (“M” version) or from sequencing summaries and *.fastq* files (“G” version). Backward compatibility with previous MinION and GridION X5 releases is also guaranteed. In addition, we provide *NanoCompare()* to allow a quick comparison of MinION and GridION X5 analyzed data. Below are described two possible workflows (also summarized in fig 1) for users wishing to analyze their MinION or GridION X5 data with NanoR.

**Fig 1.**
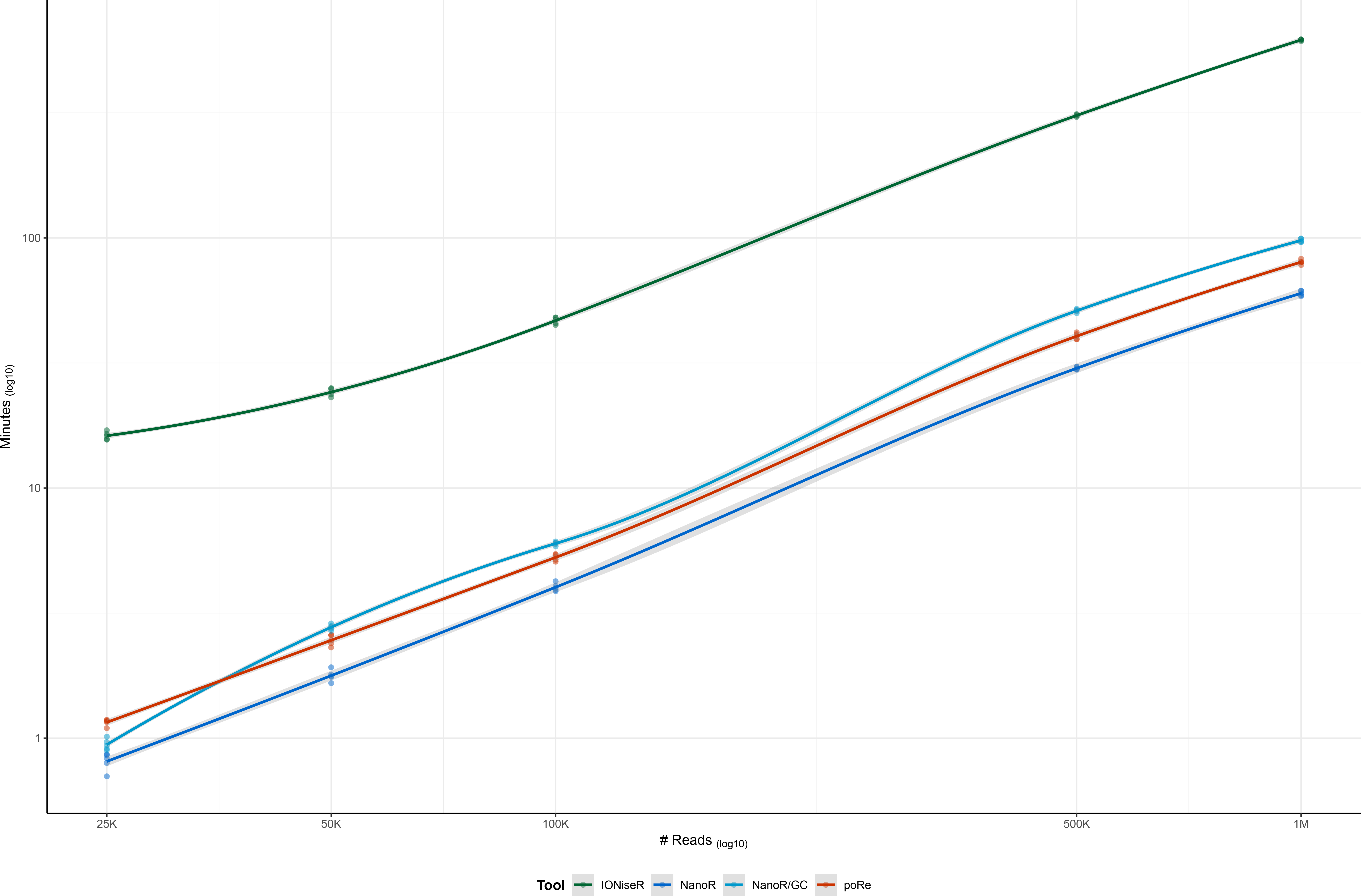
NanoR workflow. NanoR can work with both basecalled *.fast5* files and sequencing summary/*.fastq* files. Users have to rely on NanoFastqM to direclty extract.fastq sequences from basecalled .fast5 files and on NanoFastqG to filter .fastq files. NanoPrepare functions, as well as NanoTable and NanoStats can be used one after another to generate a complete overview for the sequencing run, starting from basecalled *.fast5* files (“M” version) or from sequencing summary and .*.fastq* files (“G” version). NanoCompare, at last, allows one-command comparison of MinION/GridION X5 analyzed sequencing experiments

### Basecalled *.fast5* files analysis

ONT’s MinION and GridION X5 output Guppy-basecalled *.fast5* files, which optionally can be stored into multi-read *.fast5* files (4000 read for each *.fast5* file, by default), from which key informations on the sequencing experiment can be retrived. The “M” version of NanoR can be used for this purpose.

#### Data preparation

In order to prepare MinION and GridION X5 basecalled *.fast5* files for further analyses, we provide *NanoPrepareM()*: *NanoPrepareM()* organizes the input files so that they can be easily accessed by other “M” functions from this package. The user must provide to *NanoPrepareM()* the path to passed basecalled *.fast5* reads and a label that will be used to identify the experiment, while the path to failed *.fast5* reads is optional. By default, basecalled *.fast5* files given as input are considered single-read but multi-read *.fast5* files are also supported after switching to *TRUE* the *MultiRead* parameter of this function. *NanoPrepareM()* returns an object of class list required by other functions from this package.

#### Extraction of *.fastq* and *.fasta* files

As explained in the *Introduction* section of this article, MinION and GridION X5 generate *.fastq* files for both passed and failed reads using a quality treshold of 7 (sequences with quality *geq* 7 are considere passed, the others are considered failed). NanoR offers the possibility to extract *.fastq* sequences directly from passed basecalled *.fast5* files using a user-defined treshold (which is meant to be higher then the default one) and, optionally, to convert them to *.fasta* by using *NanoFastqM()*. The *.fastq* extraction process can be accellerated using multiple cores.

#### Extraction of *.fast5* files metadata

Each basecalled *.fast5* file contains metadata such as the read identifier, the number of both channel and mux that generate the read, its generation time as well as its length and quality score; *NanoTableM()* extracts all these informations from each passed *.fast5* read, optionally calculating its GC content. As metadata extraction from *.fast5* sequences can be slow for large datasets, *NanoTableM()* can be accelerated using multiple cores. Starting from the object of class list returned by *NanoPrepareM()* and a path to a folder on which the output will be saved, *NanoTableM()* returns a metadata table with 7 columns that describes each passed *.fast5* read.

#### Plot statistics

*NanoStatsM()* is the main function of the NanoR package for the basecalled *.fast5* files data analysis; the user must provide to *NanoStatsM()* the object of class list returned by *NanoPrepareM()* and the table returned by *NanoTableM()* together with the path to the same folder specified for *NanoTableM()*. Main outputs from *NanoTableM/()* are a table containing an overview of the sequencing run (*ShortSummary.txt*), and 5 *.pdf* files representing:

- plots of cumulative reads and base pairs produced during the experiment
- plots of reads number, base pairs number, reads length (min, average and max) and reads quality (min, average and max) calculated every half an hour of experiment. (Fig 2)
- plot of read length and read quality compared jointly (Fig 3)
- plots of channels and muxes activity with respect to their real disposition on the MinION flowcell (these plots are useful to diagnose problems in particular areas of the flowcell) (Fig 4)
- plots that summarize number and percentage of passed and failed/skipped *.fast5* reads as well as GC content count (if computed by *NanoTableM()*) calculated for passed ones

**Fig 2.**
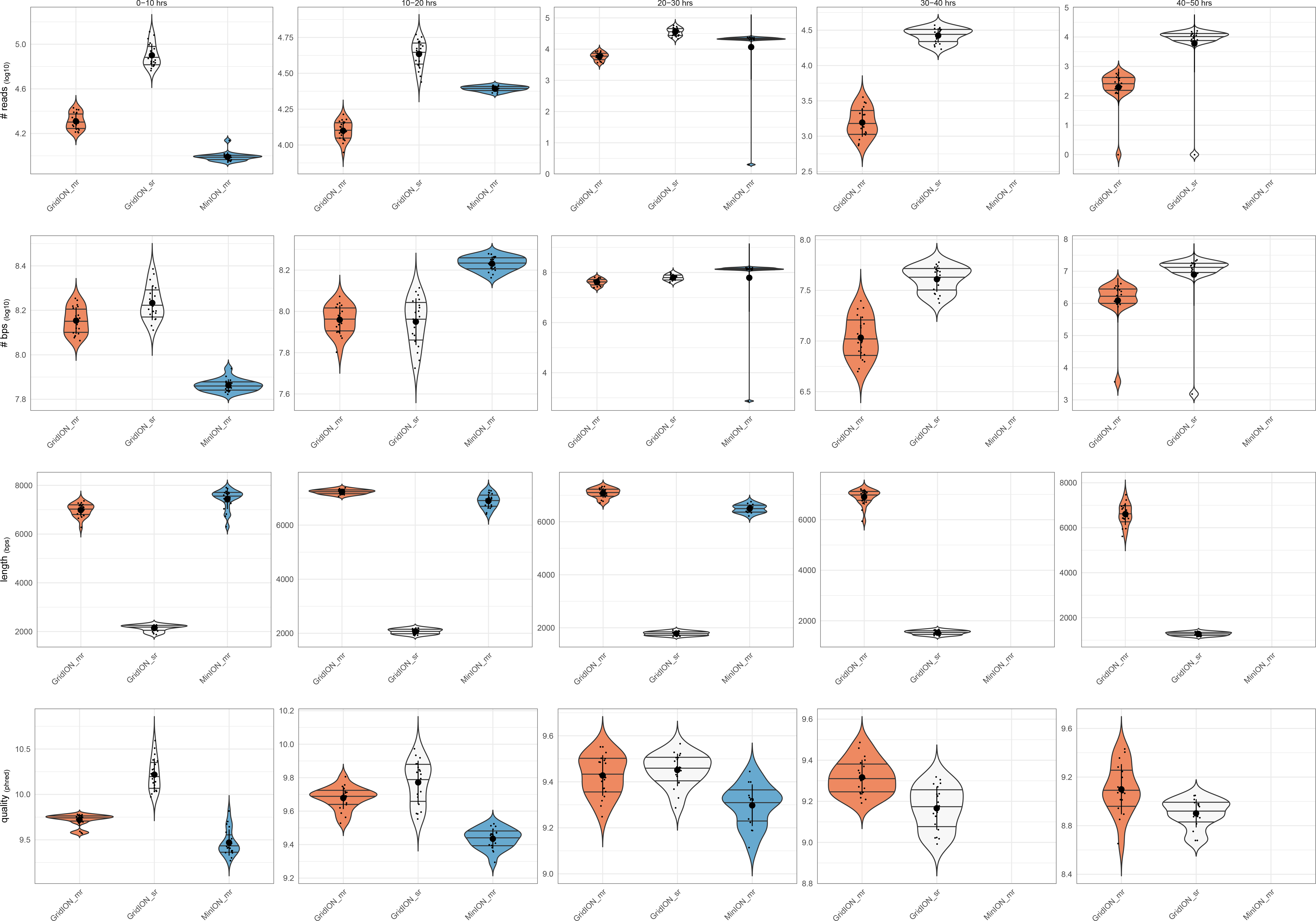
Reads number, base pairs number, reads length and reads quality per-time bin. The plots show the number of reads (**Panel A**) and basepairs (**Panel B**), the maximum, average and minimum length of reads in log10 scale (**Panel C**) and the maximum, average and minimum quality of reads (**Panel D**), all calculated every 30 minutes of an experimental MinION run

**Fig 3.**
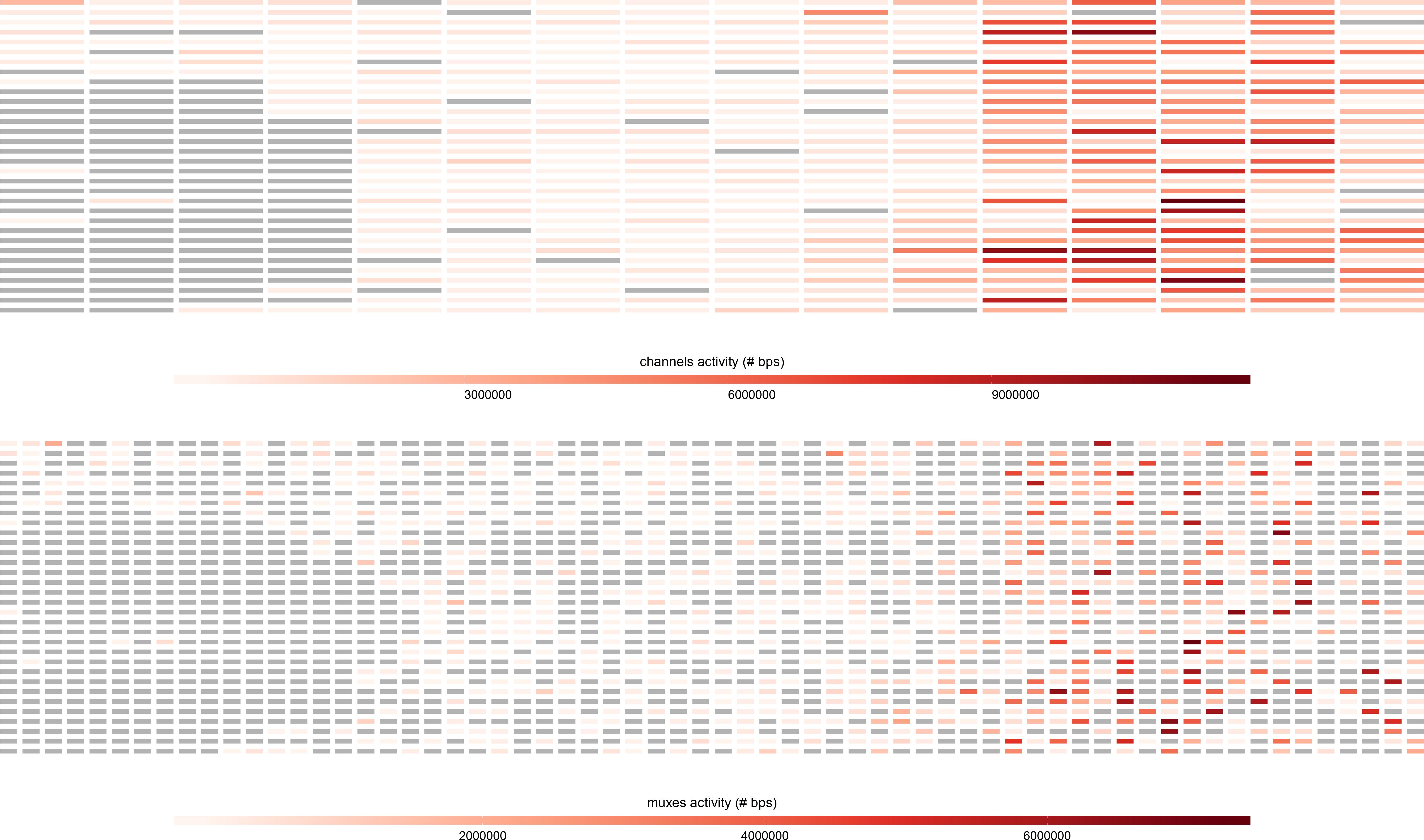
Reads length and reads quality compared jointly. The plot shows the correlation between length (x axis) and quality (y axis) for ~ 100000 MinION reads. The regression line highlights that longer the reads are, higher their quality score is.

**Fig 4.**
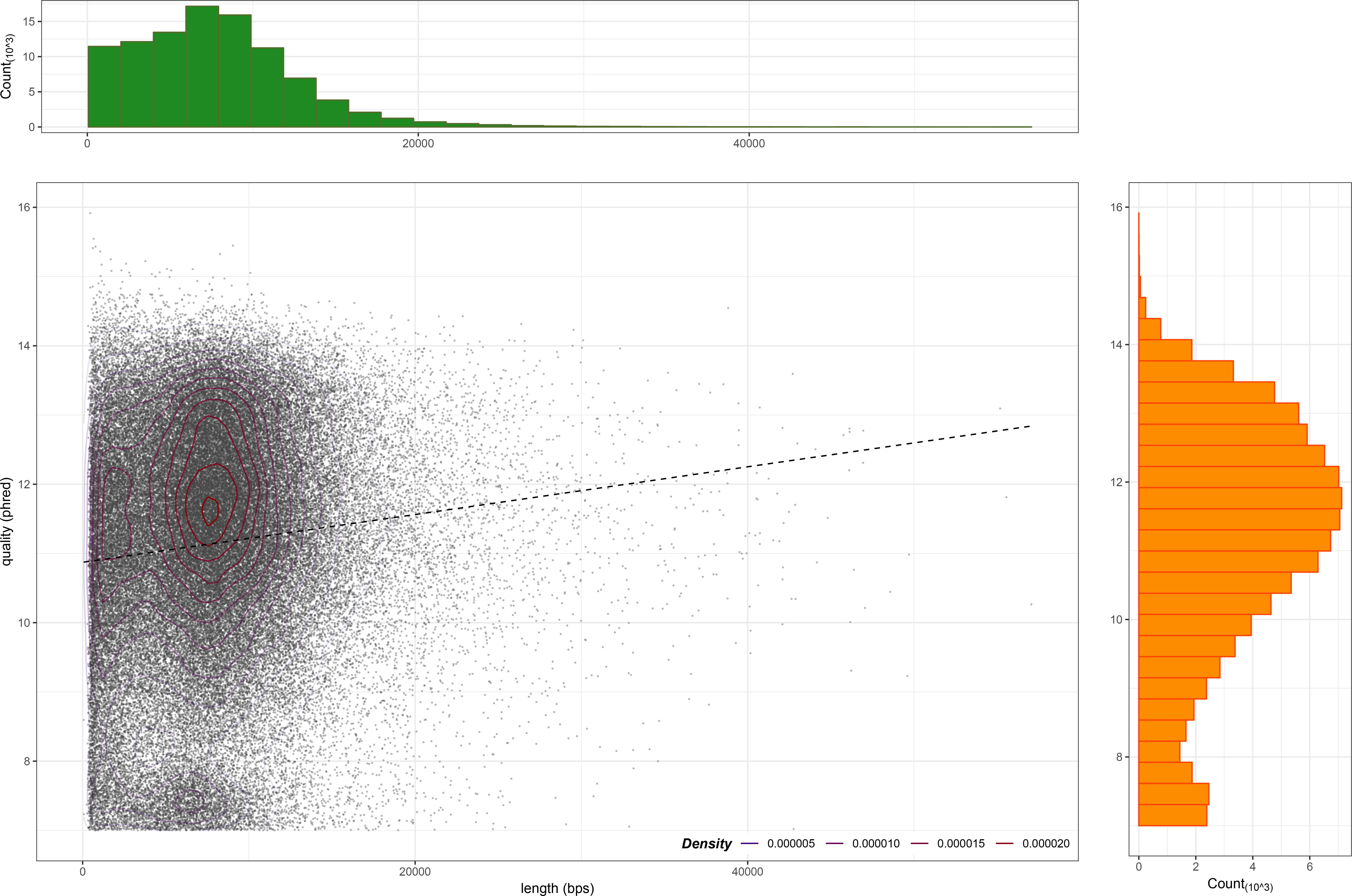
Heatmap of channels and muxes activity. The plots show the base pairs productivity of channels (**Panel A**) and muxes (**Panel B**) with respect to their real disposition on the Flow Cell (as described in https://community.nanoporetech.com/technical_documents/hardware/v/hwtd_5000_v1_revh_03may2016/flow-cell-chip) for a MinION run; inactive channels and muxes are grey-colored.

### Sequencing summary and *.fastq* files analysis

Together with basecalled *.fast5* files, MinION and GridION X5 output a single sequencing summary file which contains metadata that describe each sequenced read. This sequencing summary file can be used, together with *.fastq* files, as input to generate a complete overview for the experimental run. The “G” version of NanoR is built on 4 functions specular to those described for the “M” version. Briefly, *NanoPrepareG()* organizes the sequencing summary and *.fastq* files so that they can be easily accessed by other “G” functions from this package, *NanoTableG()* retain the most useful informations from the sequencing summary file and, optionally, calculate GC contentent for passed *.fastq*; *NanoStatG()* plots statistics and *FastqFilterG()* filters passed *.fastq* files using a user-defined treshold, optionally converting them to *.fasta* as well.

### Compare MinION/GridION X5 data

When dealing with 2 or more MinION/GridION X5 experiments, users may wish to compare their sequencing runs: to this purpose we provide *NanoCompare()*. *NanoCompare()* takes, as inputs, a vector containing the paths to the *DataForComparison* folders generated by *NanoStatsM()*/*NanoStatsG()*, an ordered vector contaning identifiers to name each experiment and the path to the folder in which *NanoCompare()* plots will be saved. *NanoCompare()* returns a *.pdf* file containing violin plots (Fig 5) that compare reads number, base pairs number, reads length and reads quality for the given experiments.

**Fig 5.**
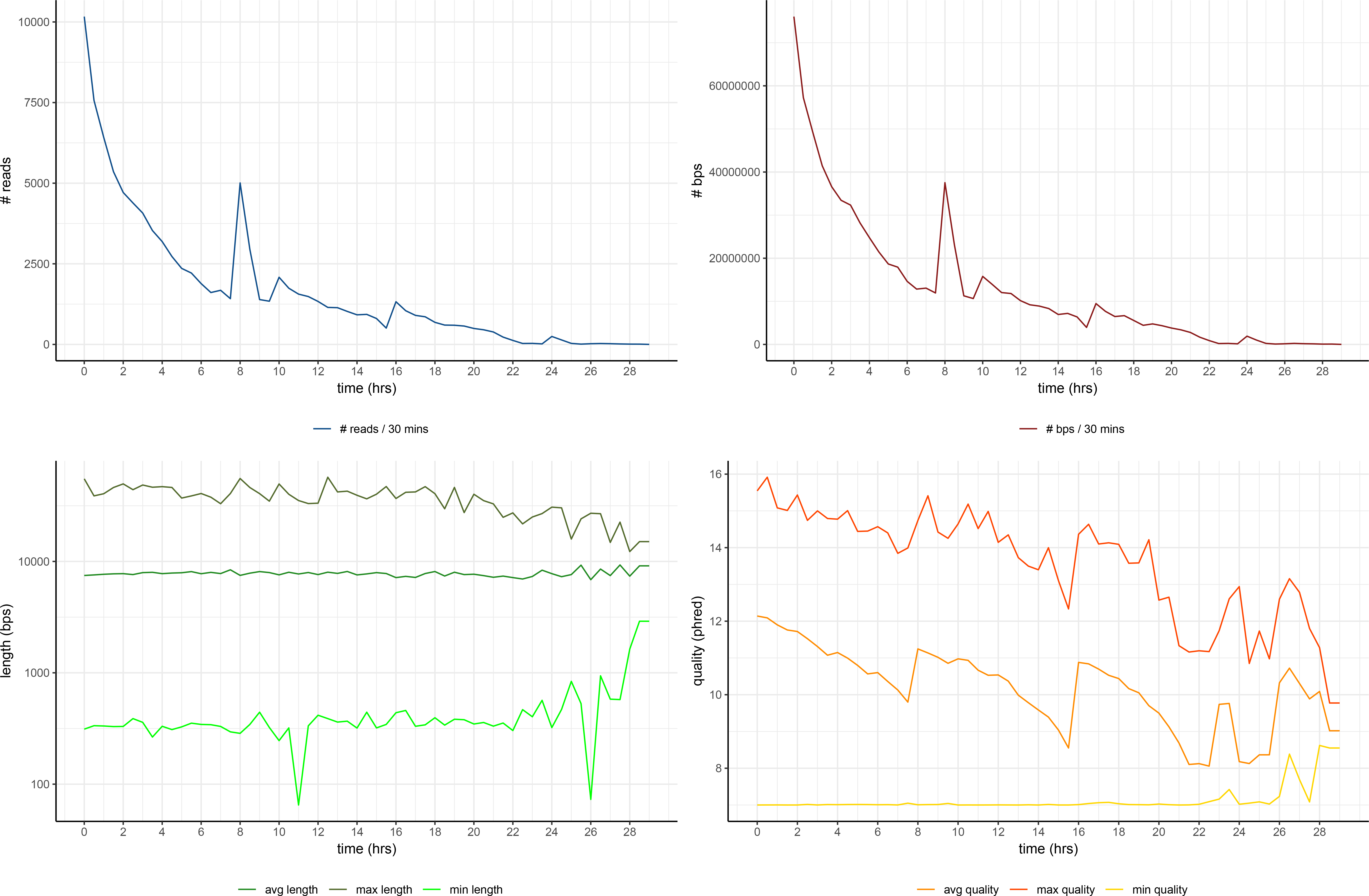
Violin plots for comparison between experiments. From top to bottom, the plots show the comparison 3 ONT experiments (first 2 are GridION X5 experiments, producin multi-read and single-read *.fast5* files respectively and last is a MinION experiment producing multi-read *.fast5* files) in term of reads number, base pairs number, reads mean length and reads mean quality. Comparison is done every 10 hours of experiment using time bins of 30 minutes.

## Discussion

As described in detail in the previous section, NanoR is based on very few functions (4 for basecalled data anaysis, 4 for sequencing summary and *.fastq* files analysis and 1 to compare MinION/GridION x5 experiments), thus being intuitive and easy to use, even for biologists with little knowledge of R language. It also produces meaningful data within acceptable time frames. Indeed, the most time-consuming step when dealing with basecalled *.fast5* files is the metadata extraction. In order to demonstrate how our tool performs compared to other tools that run in the same environment, we compared the user waiting-time when extracting metadata from single-read MinION basecalled *.fast5* files with 3 different R packages: the NanoR package (using *NanoTableM()*), with and without GC content calculation), the poRe package (using *extract.fast5()*, taken from the poRe parallel GUI and used to extract only metadata (*i.e.* the read filename, the channel number, the read number, the read start time, the length of the sequence, the run identifier, the read identifier, the experiment start time and a logical expression indicating whether the read is passed (T) or not (F).) and the IONiseR [17] package (using *readFast5Summary()* and *readInfo()* (combined, these functions from IONiseR returns the filename, the number of both channel and mux that generate a certain sequence and its pass/fail status). In particular we randomly sampled increasing number of reads (25000,50000,100000,500000,1000000) from a total of ~ 2000000 reads taken from 5 different MinION sequencing runs and we calculated the user waiting-time when extracting metadata from them. The aforementioned functions were parallelized with the R “parallel” package using 10 Intel®Xeon®CPU E5-46100 @ 2.40GHz cores on a 48 cores SUSE Linux Enterprise Server 11.

For each number of reads analyzed, we repeated the sampling-extraction step 5 times (all the time measures, together with their averages and standard deviations are attached in S1 Supporting Information). Cache was cleaned after each metadata extraction.

Fig 6 shows these results as LOESS curves: as expected, the user waiting-time increases, for all the three packages, when increases the number of the reads analyzed, with IONiseR that turned out to be the slowest and NanoR the fastest, if it doesn’t have to read the *.fastq* information to compute GC content.

**Fig 6.**
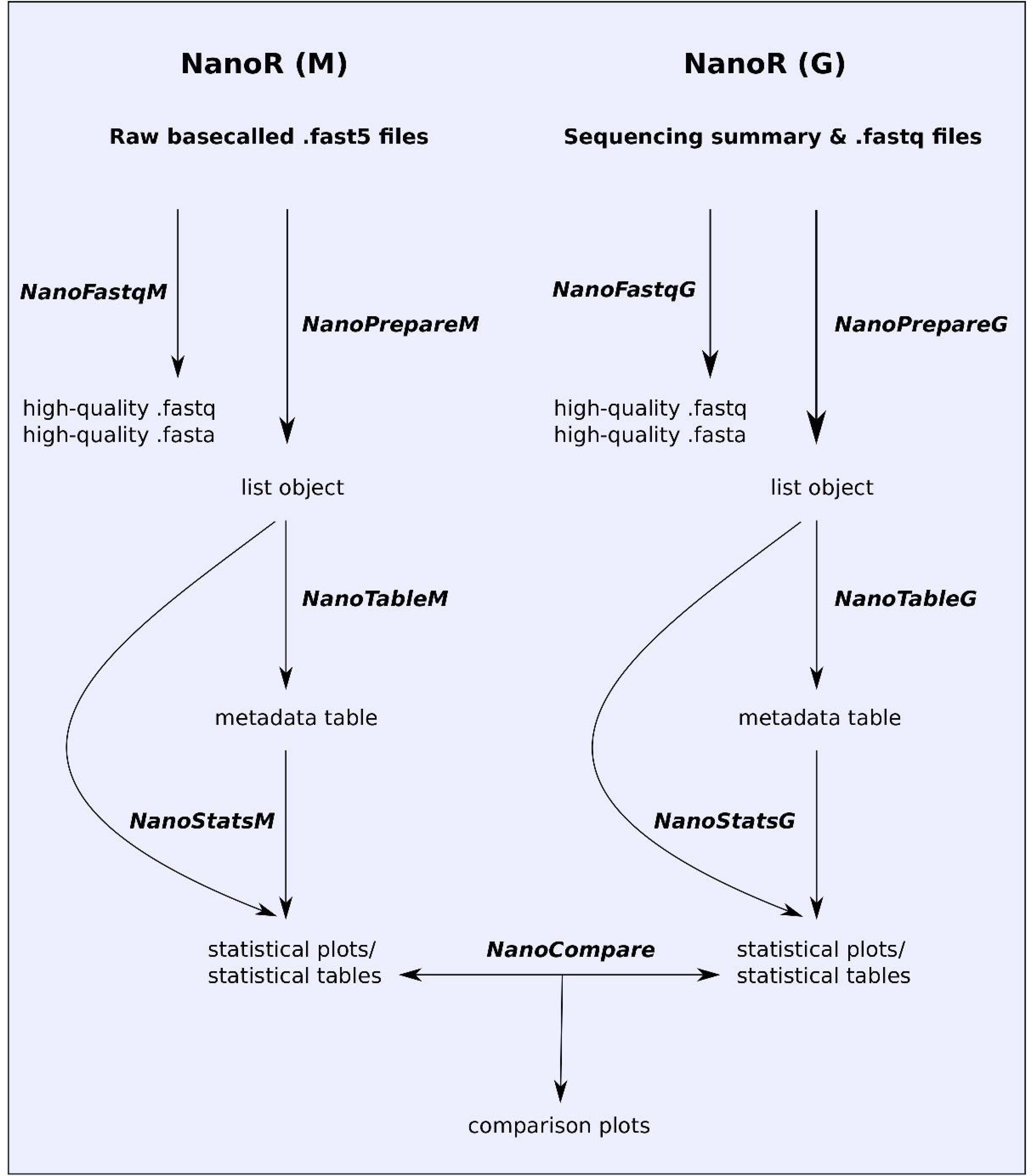
LOESS curves comparing NanoR (dark blue, light blue), poRe (red) and IONiseR (green) performances when extracting metadata informations from increasing number of *.fast5* reads (25000,50000,100000,500000,1000000) randomly sanpled from ~ 2000000 reads coming from 5 MinION runs, using 10 Intel®Xeon®CPU E5-46100 @ 2.40GHz cores. Each sampling-extraction step was repeated 5 times. Under the same conditions (*i.e.* without GC content calculation), NanoR is the fastest in extracting metadata informations for all the groups of *.fast5* file considered (*e.g.* NanoR takes approximately 60 minutes to extract metadata from 1000000 *.fast5* files, poRe takes approximately 80 minutes and IONiseR takes over 10 hours) while the extraction of metadata together with GC content computation (light blue line) makes NanoR working slightly slower (it takes approximatley 100 minutes to both extract metadata informations from 1000000 *.fast5* files and calculate their GC content). Data on x and y axes are log10-scaled.

Indeed, *NanoTable()* function, instead of dumping *.fast5* files, which requires to stream each file entirely, makes use of pointers, which allow direct access to that part of the file structure that contains the required information. This behaviour, also known as “lazy evaluation principle”, allows users to save time when retrieving informations from files with nested architectures, as it avoids to stream the entire file before fetching the desired informations. Moreover, NanoR pre-allocates a portion of computer memory to store informations from each read before the extraction and fills in values as it goes, making it fast to save retrived metadata. We have also assessed the user waiting-time when retrieving metadata from reads of different size, as the length-independent speed of analysis can be an useful feature when dealing with ultra-long reads [18]; in this case, all the three tools perform equally, as there isn’t, as expected, an increase of the user-waiting time when the size of the reads increases (except for NanoR if it had to read the *.fastq* information to compute GC content: in that case, the user-waiting time increases in a length-dependent manner). NanoR has a number of features in common with other tools built to deal with MinION sequencing data such as poRe, IONiseR, poretools and HPG pore. Table 1 summarizes NanoR features and compares them to those implemented in the aformentioned tools. As shown in Table 1, NanoR reproduces most of the output generated by the other tools for MinION single-read data analysis, overcoming their limitations. In particular, it extends the statistical analysis carried out by all the other tools (*e.g.*, it allows users to check the trend of the sequencing run every 30-minutes (Fig 2) and to compare reads length and reads quality jointly (Fig 3) as exclusive features) and is also faster then competitors in the R environment. Moreover, NanoR allows easy parallelization without the need to rely on distributed clusters, as it strictly depends for that on the R base package “parallel”. As other unique features, NanoR is, at the moment, the only tool capable to deal with outputs coming from all MinION and GridION X5 releases and allows quick comparison of MinION/GridION X5 analyzed experiments, making it easy to visualize differences between experimental sequencing runs. An example of application of NanoR on GridION X5 data is shown in S2 Supporting Information.

**Table 1.**
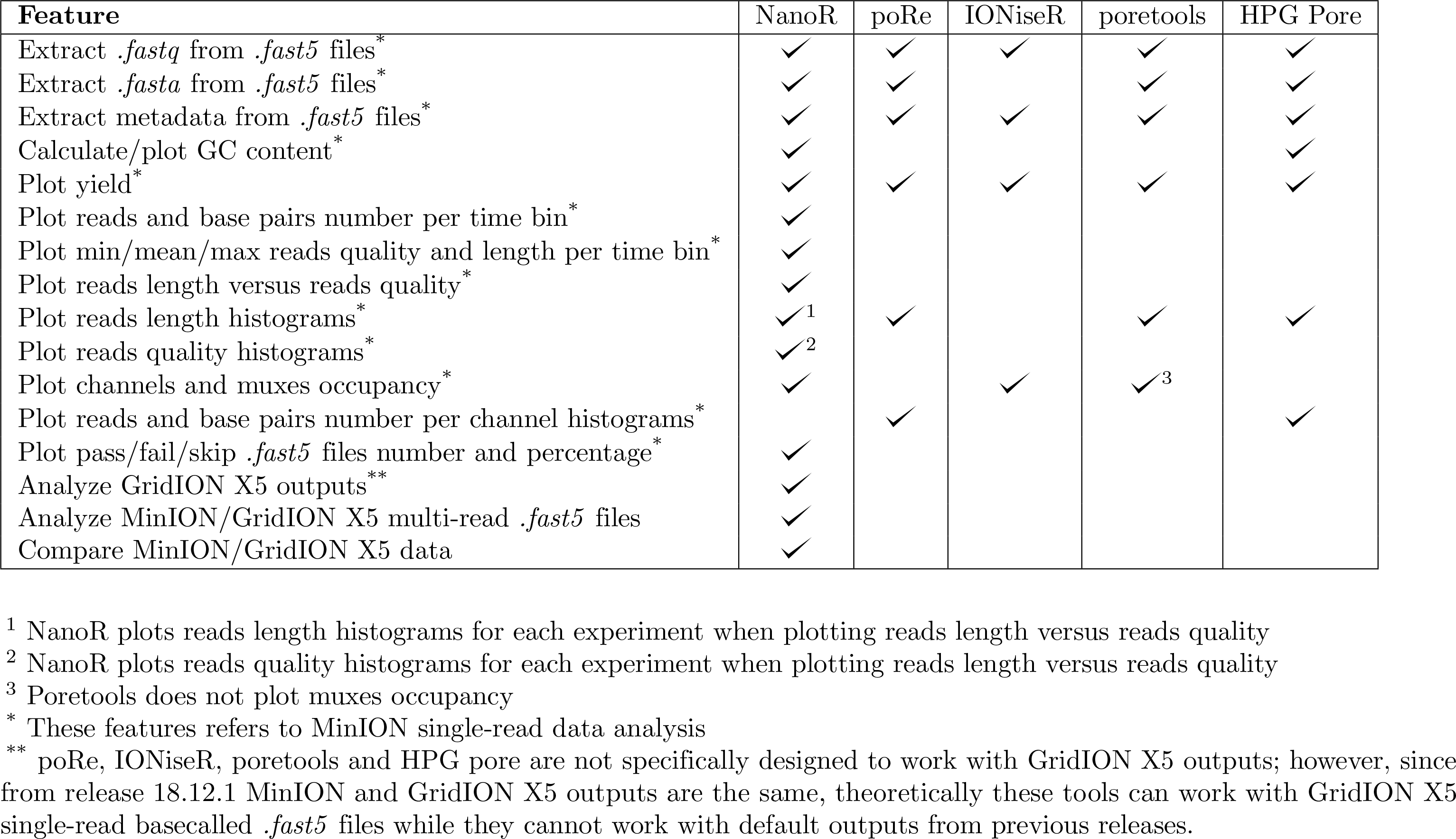
Comparison between NanoR, poRe, IONiseR, poretools and HPG pore

## Conclusion

MinION and GridION X5 from ONT are real-time sequencing devices capable to generate very long reads required for precise characterisation of complex structural variants such as copy number alterations. While library preparation is not the “hard part” when working with these devices, researchers may have some difficulties in analyzing their outputted data. Here we presented NanoR, a cross-platform package for the widespread statistical language R, that allows easy and fast analysis of MinION and GridION X5 outputs. Currently, NanoR has been extensively tested on the following 1D sequencing data:

- MinION and GridION X5 basecalled *.fast5* files (from any release till MinION 18.2 and GridION 18.2.1)
- MinION and GridION X5 sequencing summary and *.fastq* files (from any release till MinION 18.2 and GridION 18.2.1)

NanoR can be easily improved in the future in order to be up-to-date with the expected releases of MinION and GridION X5 softwares and to extend its data analysis capability and, even though there are other programs that can be used to analyze MinION data such as poRe, IONiseR, poretools and HPG pore, NanoR has unique features that make this tool extremely useful for researchers that work with ONT’s sequencing data.

## Software availability and requirements

- Project name: NanoR
- Project home page: https://github.com/davidebolo1993/NanoR
- Operating systems: Linux, Windows, MacOSX
- Programming language: R
- Other requirements: *R* ≥ 3.1.3, *ggplot*2 ≥ 2.2.1, *reshape*2 ≥ 1.4.3, *RColorBrewer* ≥ 1.1.2, *scales* ≥ 0.5.0, *gridExtra* ≥ 2.3, *ShortRead* ≥ 1.24.0, *rhdf* 5 ≥ 2.14
- License: GPLv3

A dataset (~ 11 GB) of MinION single-read basecalled *.fast5* files supporting the figures of this article together with a dataset (~ 17 GB) coming from a GridION X5 run supporting data shown in S2 Supporting Information can be found at https://faspex.embl.de/aspera/faspex/external_deliveries/611?passcode=6f668c8921cea5cc30d0a6c17ec228f6ba75936c&expiration=MjAxOS0wMi0yMlQxMDo1Njo0OVo=. The aforementioned dataset contains MinION and GridION X5 runs performed before MinION 18.2 and GridION X5 18.2.1 releases. The complete set of plots returned by NanoR as well as exhaustive installation istructions and examples of how to run NanoR functions and a folder containing all the functions on which NanoR is built can be found at https://github.com/davidebolo1993/NanoR, together with the manual of the tool.

## Supporting information

S1 Supporting Informations

S2 Supporting Informations

## Declarations

### Acknowledgements

This work has been supported by Ministero della Salute (GR-2011-02352026), and by Associazione Italiana per la Ricera sul Cancro (Investigator Grant 20307).

### Author’s contributions

Davide Bolognini wrote and tested the R code. Niccolò Bartalucci and Alessandra Mingrino prepared MinION and GridION X5 libraries. Alberto Magi and Alessandro Maria Vannucchi supervised the project. Davide Bolognini and Alberto Magi wrote the manuscript.

