## Supplementary material for "NanoR: a user-friendly R package to analyze and compare nanopore sequencing data": S1 Supporting Informations

The following procedure is meant to be an example of application of NanoR on GridION X5 sequencing outputs. A GridION X5 dataset generated before 18.12.1 release is available to be downloaded as a test at `https://faspex.embl.de/aspera/faspex/external_deliveries/611?passcode=6f668c8921cea5cc30d0a6c17ec228f6ba75936c&expiration=MjAxOS0wMi0yMlQxMDo1Njo0OVo=`. You will be likely required to install the aspera plugin in order to download the test set. Once you have downloaded and extracted the plugin, you can install it (for example, on a linux machine, just launch the shell script using the bash interpreter). Refresh the web page and TestSet folder is available to be downloaded.

Once the TestSet folder has been downloaded, *.fastq* and sequencing summary files can be extracted from TestSet/GridION.tar.

```
cd TestSet
tar -xvf GridION.tar
```

This will create a GridION folder contaning *.fastq* files and sequencing summary files for the GridION X5 run.

We can create a unique, unfiltered, *.fastq* file for further comparison from *.fastq* files in GridION folder.

```
cd TestSet && mkdir test
cat GridION/*.fastq > test/unfiltered.fq
```

Also *NanoFastqG()* can be used on *.fastq* files in sample folder. From an R console, type:

```
library(NanoR)
Data <- DataSummary <- DataFastq <- 'Path/To/TestSet/GridION'
DataOut<-'Path/To/TestSet/test'
Label<-'filtered'
NanoFastqG(Data, Data, DataOut, Cores=4, Label=Label, Minquality=10)
```

This will create in TestSet/test a folder named "filtered" with a filtered.fq file inside that we can move to the previous folder.

```
cd TestSet/test/filtered && mv filtered.fq ../.
cd .. && rm -r filtered
```

Now we we can apply a standard alignment procedure using minimap2 [1] and samtools [2].

```
cd TestSet/test
MyFiles="unfiltered filtered"
MyGenome="path/to/referencegenome" #we used HG19 for our test
for files in $MyFiles; do
minimap2 -ax map-ont -t 5 $MyGenome $files".fq" > $files".sam"
htsbox samview -bS $files".sam" > $files".bam"
rm $files".sam"
samtools sort -@ 5 $files".bam" > $files".srt.bam"
rm $files".bam"
```

```
samtools index $files".srt.bam"
done
```

Statistics on the two .bam files can be calculated using Alfred [3].

```
MyGenome='path/to/referencegenome' #we used HG19 for our test
unfiltered="unfiltered.srt.bam"
filtered="filtered.srt.bam"
alfred qc -r $MyGenome -s -u -o qc_unfiltered.tsv.gz $unfiltered
alfred qc -r $MyGenome -s -u -o qc_filtered.tsv.gz $filtered
```

Finally, we can extract the results:

```
zgrep ^ME qc_unfiltered.tsv.gz |\
cut -f 2- |\
datamash transpose |\
column -t > unfiltered.tsv

zgrep ^ME qc_filtered.tsv.gz |\
cut -f 2- |\
datamash transpose |\
column -t > filtered.tsv
```

In Table 1, unfiltered.tsv and filtered.tsv where put side-by-side. *NanoFastqG()*, by filtering out all reads with quality lower than 10, makes of course all the files for downstream analyses lighter and the analyses itself faster (Table 2). The fraction of unmapped reads drastically decreases and, more importantly, the filtering step operated by *NanoFasqtG()* reduces the Error Rate of the alignment file (unfiltered.srt.bam has an error rate of .12 while filtered.srt.bam an error rate of .09), which is notorously a crucial problem for accurate detection of structural variants.

Table 1: Statistics for bamfiles

| Sample | unfiltered.srt | filtered.srt |
|---|---|---|
| Library | DefaultLib | DefaultLib |
| #QCFail | 0 | 0 |
| QCFailFraction | 0 | 0 |
| #DuplicateMarked | 0 | 0 |
| DuplicateFraction | 0 | 0 |
| #Unmapped | 389109 | 6782 |
| UnmappedFraction | 0.0664869 | 0.00288626 |
| #Mapped | 5463307 | 2342970 |
| MappedFraction | 0.933513 | 0.997114 |
| #MappedRead1 | 5463307 | 2342970 |
| #MappedRead2 | 0 | 0 |
| RatioMapped2vsMapped1 | 0 | 0 |
| #MappedForward | 2733801 | 1169518 |
| MappedForwardFraction | 0.500393 | 0.49916 |
| #MappedReverse | 2729506 | 1173452 |
| MappedReverseFraction | 0.499607 | 0.50084 |
| #SecondaryAlignments | 1361238 | 432206 |
| SecondaryAlignmentFraction | 0.24916 | 0.184469 |
| #SupplementaryAlignments | 98568 | 43012 |
| SupplementaryAlignmentFraction | 0.0180418 | 0.0183579 |
| #SplicedAlignments | 0 | 0 |
| SplicedAlignmentFraction | 0 | 0 |
| #Pairs | 0 | 0 |
| #MappedPairs | 0 | 0 |
| MappedPairsFraction | 0 | 0 |
| #MappedSameChr | 0 | 0 |
| MappedSameChrFraction | 0 | 0 |
| #MappedProperPair | 0 | 0 |
| MappedProperFraction | 0 | 0 |

| Sample | unfiltered.minimap2.srt | filtered.minimap2.srt |
|---|---|---|
| #ReferenceBp | 3137161264 | 3137161264 |
| #ReferenceNs | 239850802 | 239850802 |
| #AlignedBases | 7105376143 | 3700886273 |
| #MatchedBases | 6780206420 | 3584535165 |
| MatchRate | 0.954236 | 0.968561 |
| #MismatchedBases | 325169723 | 116351108 |
| MismatchRate | 0.0457639 | 0.0314387 |
| #DeletionsCigarD | 232610065 | 91561199 |
| DeletionRate | 0.0327372 | 0.0247403 |
| HomopolymerContextDel | 0.343239 | 0.377105 |
| #InsertionsCigarI | 306013346 | 123375461 |
| InsertionRate | 0.0430679 | 0.0333367 |
| HomopolymerContextIns | 0.3793 | 0.416704 |
| #SoftClippedBases | 10654263 | 4563181 |
| SoftClipRate | 0.00149947 | 0.001233 |
| #HardClippedBases | 196512 | 85645 |
| HardClipRate | 2.76568e-05 | 2.31418e-05 |
| ErrorRate | 0.123096 | 0.0907719 |
| MedianReadLength | 1271 | 1398 |
| DefaultLibraryLayout | 0 | 0 |
| MedianInsertSize | 0 | 0 |
| MedianCoverage | 2 | 1 |
| SDCoverage | 25.3881 | 14.212 |
| CoveredBp | 2568074701 | 1988707221 |
| FractionCovered | 0.886365 | 0.686398 |
| BpCov1ToCovNRatio | 0.244871 | 0.468283 |
| BpCov1ToCov2Ratio | 0.902406 | 1.57344 |
| MedianMAPQ | 60 | 60 |

Table 2 compares real times, in seconds, when aligning unfiltered.fq and filtered.fq files with minimap2, using 5 Intel®Xeon®CPU E5-46100 @ 2.40GHz cores on a 48 cores SUSE Linux Enterprise Server 11. The alignment step was repeated 5 times for each *.fastq* file.

Table 2: Time comparison for alignment

| #Replicates | unfiltered.fq (seconds) | filtered.fq (seconds) |
|---|---|---|
| 1 | 4052.23 | 1556.84 |
| 2 | 3973.41 | 1544.03 |
| 3 | 3942.55 | 1557.75 |
| 4 | 3984.53 | 1474.11 |
| 5 | 3854.45 | 1442.74 |

For unfiltered.fq mean is 3961.434 seconds and sd 1556.84 seconds, while for filtered.fq mean is 1515.094 seconds and sd 71.97733 seconds.
