## Supplementary material for "NanoR: a user-friendly R package to analyze and compare nanopore sequencing data": S2 Supporting Informations

Table 1 describes time replicates for each tool. Tools are NanoR (without GC content calculation - NanoR/NOGC - and with GC content calculation - NanoR/GC -), poRE (without GC content calculation - poRe/NOGC -) and IONiseR (without GC content calculation - IONiseR/NOGC - ). # Reads is expressed as log10 value. Time is expressed in minutes. We sampled 5 times for each number of reads.

Table 1: Time replicates

| Tool | # Reads (log10) | Time (minutes) | # Replicate |
|---|---|---|---|
| NanoR/NOGC | 4.39794000867204 | 0.795823080937068 | 1 |
| NanoR/NOGC | 4.39794000867204 | 0.862223080937068 | 2 |
| NanoR/NOGC | 4.39794000867204 | 0.855823080937068 | 3 |
| NanoR/NOGC | 4.39794000867204 | 0.703423080937068 | 4 |
| NanoR/NOGC | 4.39794000867204 | 0.829323080937068 | 5 |
| NanoR/NOGC | 4.69897000433602 | 1.75740079085032 | 1 |
| NanoR/NOGC | 4.69897000433602 | 1.79600079085032 | 2 |
| NanoR/NOGC | 4.69897000433602 | 1.76770079085032 | 3 |
| NanoR/NOGC | 4.69897000433602 | 1.91950079085032 | 4 |
| NanoR/NOGC | 4.69897000433602 | 1.65940079085032 | 5 |
| NanoR/NOGC | 5 | 3.99064147988955 | 1 |
| NanoR/NOGC | 5 | 4.06204147988955 | 2 |
| NanoR/NOGC | 5 | 3.91734147988955 | 3 |
| NanoR/NOGC | 5 | 3.86544147988955 | 4 |
| NanoR/NOGC | 5 | 4.23814147988955 | 5 |
| NanoR/NOGC | 5.69897000433602 | 29.7442908044656 | 1 |
| NanoR/NOGC | 5.69897000433602 | 29.8996908044656 | 2 |
| NanoR/NOGC | 5.69897000433602 | 30.5630908044656 | 3 |
| NanoR/NOGC | 5.69897000433602 | 30.5420908044656 | 4 |
| NanoR/NOGC | 5.69897000433602 | 29.8750908044656 | 5 |
| NanoR/NOGC | 6 | 59.9819412811597 | 1 |
| NanoR/NOGC | 6 | 58.5662412811597 | 2 |
| NanoR/NOGC | 6 | 59.0579412811597 | 3 |
| NanoR/NOGC | 6 | 61.3805412811597 | 4 |
| NanoR/NOGC | 6 | 61.4781412811597 | 5 |
| poRe/NOGC | 4.39794000867204 | 1.0964647491773 | 1 |
| poRe/NOGC | 4.39794000867204 | 1.1679647491773 | 2 |
| poRe/NOGC | 4.39794000867204 | 1.1600647491773 | 3 |
| poRe/NOGC | 4.39794000867204 | 1.1831647491773 | 4 |
| poRe/NOGC | 4.39794000867204 | 1.1774647491773 | 5 |
| poRe/NOGC | 4.69897000433602 | 2.39600941737493 | 1 |
| poRe/NOGC | 4.69897000433602 | 2.58090941737493 | 2 |
| poRe/NOGC | 4.69897000433602 | 2.30480941737493 | 3 |
| poRe/NOGC | 4.69897000433602 | 2.46100941737493 | 4 |
| poRe/NOGC | 4.69897000433602 | 2.58220941737493 | 5 |
| poRe/NOGC | 5 | 5.16840459704399 | 1 |
| poRe/NOGC | 5 | 5.40890459704399 | 2 |
| poRe/NOGC | 5 | 5.07890459704399 | 3 |
| poRe/NOGC | 5 | 5.44230459704399 | 4 |
| poRe/NOGC | 5 | 5.30920459704399 | 5 |
| poRe/NOGC | 5.69897000433602 | 40.5475916838646 | 1 |
| poRe/NOGC | 5.69897000433602 | 41.2082916838646 | 2 |
| poRe/NOGC | 5.69897000433602 | 39.4399916838646 | 3 |
| poRe/NOGC | 5.69897000433602 | 41.9332916838646 | 4 |
| poRe/NOGC | 5.69897000433602 | 39.2783916838646 | 5 |
| poRe/NOGC | 6 | 80.318941227595 | 1 |
| poRe/NOGC | 6 | 79.809741227595 | 2 |
| poRe/NOGC | 6 | 77.977141227595 | 3 |
| poRe/NOGC | 6 | 82.437241227595 | 4 |
| poRe/NOGC | 6 | 79.876941227595 | 5 |

| Tool | # Reads (log10) | Time (minutes) | # Replicate |
|---|---|---|---|
| IONiseR/NOGC | 4.39794000867204 | 16.4624835574627 | 1 |
| IONiseR/NOGC | 4.39794000867204 | 15.6150835574627 | 2 |
| IONiseR/NOGC | 4.39794000867204 | 17.0218835574627 | 3 |
| IONiseR/NOGC | 4.39794000867204 | 16.1810835574627 | 4 |
| IONiseR/NOGC | 4.39794000867204 | 15.6802835574627 | 5 |
| IONiseR/NOGC | 4.69897000433602 | 23.6942521437009 | 1 |
| IONiseR/NOGC | 4.69897000433602 | 24.2224521437009 | 2 |
| IONiseR/NOGC | 4.69897000433602 | 23.0433521437009 | 3 |
| IONiseR/NOGC | 4.69897000433602 | 25.0779521437009 | 4 |
| IONiseR/NOGC | 4.69897000433602 | 24.9036521437009 | 5 |
| IONiseR/NOGC | 5 | 44.890476851066 | 1 |
| IONiseR/NOGC | 5 | 45.587576851066 | 2 |
| IONiseR/NOGC | 5 | 47.111276851066 | 3 |
| IONiseR/NOGC | 5 | 48.022376851066 | 4 |
| IONiseR/NOGC | 5 | 48.094176851066 | 5 |
| IONiseR/NOGC | 5.69897000433602 | 308.592011109988 | 1 |
| IONiseR/NOGC | 5.69897000433602 | 310.297511109988 | 2 |
| IONiseR/NOGC | 5.69897000433602 | 312.448411109988 | 3 |
| IONiseR/NOGC | 5.69897000433602 | 304.563911109988 | 4 |
| IONiseR/NOGC | 5.69897000433602 | 308.288911109988 | 5 |
| IONiseR/NOGC | 6 | 621.416425133149 | 1 |
| IONiseR/NOGC | 6 | 622.511825133149 | 2 |
| IONiseR/NOGC | 6 | 612.641525133149 | 3 |
| IONiseR/NOGC | 6 | 621.115025133149 | 4 |
| IONiseR/NOGC | 6 | 617.907225133149 | 5 |
| NanoR/GC | 4.39794000867204 | 0.93423056046168 | 1 |
| NanoR/GC | 4.39794000867204 | 0.96503056046168 | 2 |
| NanoR/GC | 4.39794000867204 | 0.90703056046168 | 3 |
| NanoR/GC | 4.39794000867204 | 1.01403056046168 | 4 |
| NanoR/GC | 4.39794000867204 | 0.89493056046168 | 5 |
| NanoR/GC | 4.69897000433602 | 2.73793083349864 | 1 |
| NanoR/GC | 4.69897000433602 | 2.80623083349864 | 2 |
| NanoR/GC | 4.69897000433602 | 2.87443083349864 | 3 |
| NanoR/GC | 4.69897000433602 | 2.77803083349864 | 4 |
| NanoR/GC | 4.69897000433602 | 2.68713083349864 | 5 |
| NanoR/GC | 5 | 5.97110960205396 | 1 |
| NanoR/GC | 5 | 6.02810960205396 | 2 |
| NanoR/GC | 5 | 6.04890960205396 | 3 |
| NanoR/GC | 5 | 5.84370960205396 | 4 |
| NanoR/GC | 5 | 6.11400960205396 | 5 |
| NanoR/GC | 5.69897000433602 | 50.9489521197478 | 1 |
| NanoR/GC | 5.69897000433602 | 52.0769521197478 | 2 |
| NanoR/GC | 5.69897000433602 | 51.4598521197478 | 3 |
| NanoR/GC | 5.69897000433602 | 49.9982521197478 | 4 |
| NanoR/GC | 5.69897000433602 | 51.2334521197478 | 5 |
| NanoR/GC | 6 | 97.3388202206294 | 1 |
| NanoR/GC | 6 | 99.3776202206294 | 2 |
| NanoR/GC | 6 | 96.5978202206294 | 3 |
| NanoR/GC | 6 | 99.4931202206294 | 4 |
| NanoR/GC | 6 | 96.0581202206294 | 5 |

Table 2 describes average time (minutes) and standard deviation (minutes) for metadata extraction for each number of reads, for each tool. Tools are NanoR (without GC content calculation - NanoR/NOGC - and with GC content calculation - NanoR/GC -), poRE (without GC content calculation - poRe/NOGC -) and IONiseR (without GC content calculation - IONiseR/NOGC - ). # Reads is expressed as log10 value.

Table 2: Time replicates: mean and sd

| Tool | # Reads(log10) | Mean time (minutes) | Sd (minutes) |
|---|---|---|---|
| NanoR/NOGC, | 4.39794000867204 | 0.809323080937068 | 0.0647258062908451 |
| NanoR/NOGC, | 4.69897000433602 | 1.78000079085032 | 0.0934380275904837 |
| NanoR/NOGC, | 5 | 4.01472147988955 | 0.145346919472 |
| NanoR/NOGC, | 5.69897000433602 | 30.1248508044656 | 0.394983073055036 |
| NanoR/NOGC, | 6 | 60.0929612811597 | 1.32202984913352 |
| poRe/NOGC | 4.39794000867204 | 1.1570247491773 | 0.0349911846041256 |
| poRe/NOGC | 4.69897000433602 | 2.46498941737493 | 0.120010091242362 |
| poRe/NOGC | 5 | 5.28154459704399 | 0.155477532138891 |
| poRe/NOGC | 5.69897000433602 | 40.4815116838646 | 1.13715682163895 |
| poRe/NOGC | 6 | 80.084001227595 | 1.59294434554381 |
| IONiseR/NOGC | 4.39794000867204 | 16.1921635574627 | 0.582381620589112 |
| IONiseR/NOGC | 4.69897000433602 | 24.1883321437009 | 0.845482830694982 |
| IONiseR/NOGC | 5 | 46.741176851066 | 1.446110291437 |
| IONiseR/NOGC | 5.69897000433602 | 308.838151109988 | 2.90667705172075 |
| IONiseR/NOGC | 6 | 619.118405133149 | 4.00623321113487 |
| NanoR/GC | 4.39794000867204 | 0.94305056046168 | 0.04798970722978 |
| NanoR/GC | 4.69897000433602 | 2.77675083349864 | 0.0706236999880352 |
| NanoR/GC | 5 | 6.00116960205396 | 0.101777934740296 |
| NanoR/GC | 5.69897000433602 | 51.1434921197478 | 0.763043984970724 |
| NanoR/GC | 6 | 97.7731002206294 | 1.58461287291249 |
